# POLLINATOR GROOMING BEHAVIOR ALTERS POLLEN LANDSCAPES ON BEES’ BODIES AND INCREASES POLLEN CARRYOVER TO OTHER FLOWERS

**DOI:** 10.1101/2022.07.14.499901

**Authors:** Vanessa Gonzaga Marcelo, Flávia Maria Darcie Marquitti, Mario Vallejo-Marín, Vinícius Lourenço Garcia de Brito

**Affiliations:** Programa de Pós Graduação em Ecologia, Conservação e Biodiversidade, Universidade Federal de Uberlândia, Uberlândia, Brazil; Instituto de Física “Gleb Wataghin”, Universidade Estadual de Campinas, Campinas, Brasil; Department of Biological and Environmental Sciences, University of Stirling, Stirling, Scotland; Instituto de Biologia, Universidade Federal de Uberlândia, Uberlândia, Brasil

**Keywords:** pollen carryover, pollen delivery, pollen fates, pollen landscape, pollen layering, pollen placement, pollen transfer, spatially explicit agent-based model

## Abstract

Pollen participates both as the carrier of male gametes in the reproduction of flowering plants and as a key resource exploited by floral visitors, especially bees. Pollinator behavior significantly alters the patterns of pollen removal and deposition, often called pollen fates. To date, few theoretical investigations have attempted to jointly model patterns of pollen transfer and pollinator behavior, and empirical studies are restricted to species to which pollen movement can be tracked. Here we use a spatially explicit agent-based modeling approach, to determine how bee grooming behavior may alter pollen fates and affect plant reproduction. Specifically, we asked whether pollen mixing and removal during pollen grooming may change the “pollen landscape” on a bee’s body consequently affecting both pollen export by the anthers and deposition onto stigmas. Our model shows that both mixing and removal behaviors restructure the “pollen landscape” on the bee’s body, increasing pollen carryover (deposition in consecutive visits), and increasing pollen diversity (number of pollen donors) onto stigmas in sequential flower visits. Our results counterintuitively show that pollen grooming may have a positive effect on both male and female finesses during plant reproduction.

## INTRODUCTION

Most flowering plants (ca. of 90%) depend on animal vectors for pollen transport and delivery (Ollerton et al. 2011; Ollerton 2017). Animal vectors increase the siring barriers that potentially diminish the probability of male success as well as the technical difficulties in empirically tracing the pollen pathway to paternity (Minnaar et al. 2019). During pollen transfer, pollen can be covered by conspecific or heterospecific pollen (pollen layering), and/or they can be lost or displaced for instance by the behavior of the pollen vector (pollen grooming) (Harder and Barrett 1996; Michener et al. 1978; Thorp 2000; Barrett 1998). Pollen layering occurs when pollen grains are buried by layers of pollen deposited on vectors’ bodies in sequential flower visits (Price and Waser 1982; Lertzman and Gass 1983). Pollen grooming occurs when pollen vectors displace pollen grains on their bodies, from the original sites of pollen placement by anthers to new sites and this action has mostly been understood as a reproductive cost for plants. Both pollen layering and grooming potentially alter the 3-D distribution and donor composition of pollen on the bodies of pollen vectors during sequential floral visits, generating a dynamic pollen landscape (Lertzman and Gass 1983; Harder and Wilson 1989; Morris et al. 1995). Pollen landscapes were first theorized 40 years ago (Lertzman 1981), but how they may be altered by successive layering and grooming and its potential impact on male and female components of plant reproductive success has been rarely studied from both empirical and theoretical perspectives.

During sequential floral visits, different pollen vector groups may vary in their interaction with pollen grains placed in their bodies. Some vectors are not expected to move pollen grains within their bodies, while other ones may perform grooming. Specifically in bees, grooming leads to the removal of pollen grains to their scopa, where pollen is no longer useful for pollination (Thorp 1979). In contrast, groomed pollen grains that are not moved to scopa can be mixed and reach body areas different from where they were initially placed (Harder and Barrett 1996). Sequential pollen placement by different flowers followed by the effects of grooming, either removal or mixing, have the potential to alter the pollen landscape within the vectors’ bodies. In fact, in theoretical multi-donor pollen landscapes, even deeply buried pollen grains might resurface on vectors’ body and still be deposited on stigmas (Lertzman 1981; Lertzman and Gass 1983; Morris et al. 1995). Although early pollen landscape models considered pollen transport only by non-grooming vectors, like hummingbirds, vector grooming behavior was later incorporated (Harder and Wilson 1989; Harder and Barrett 1996). These later models analyzed pollen dispersal by comparing the vertical structure formed by sequential placement that results in layers and the horizontal structure formed by different intensity of contact with anthers and stigmas that can be disrupted by grooming behavior (Harder and Wilson 1989). Despite this complex and dynamical spatial arrangement of individual pollen grains from different donors on vector bodies along sequential visits being hard to describe, it may generate hotspots and cold spots of outcrossed pollen, ultimately affecting male and female components of plant reproductive success (Minnaar and Anderson 2021).

Empirical studies that assess pollen transfer and delivery require tracking the movement of individual pollen grains (Minnaar et al. 2019). Given the difficulty to achieve this pollen-level resolution in practice, empirical studies are often restricted to species that present trackable pollen grains either because they are aggregated in pollinaria (Johnson and Harder 2018; Harder et al. 2021), or because they can be distinguished by size and color polymorphisms (Thomson 1986; Harder and Thomson 1989; Luo et al. 2008; Wang et al. 2018). Other studies have observed pollen transfer by staining techniques with fluorescent powders (Lertzman 1981; Price and Waser 1982; Waser and Price 1984) or quantum dots (Minnaar and Anderson 2019). However, these techniques are still difficult to apply in more detailed studies about pollen landscape formed after sequential visits of several individual flowers, mainly because material analogous to pollen may influence the results or due to the low number of colors available (Minnaar and Anderson 2019). In this sense, theoretical studies may help to understand the influence of grooming behavior and layering in the structure of pollen landscape on the vector’s body and their effect on plant reproductive success, generating hypothesis that can be further investigated by such empirical studies.

In this study, we use a spatially explicit agent-based model (ABMs) to understand the influence of grooming behavior and layering in the structure of pollen landscape in the vector’s body and their effect on plant reproductive success. We address two questions: 1) How does the pollen landscape on a vector’s body changes after several flower visits? 2) What is the potential effect of pollen grooming and layering in pollen delivery to conspecific stigmas and in the diversity of the pollen grains received? We simulated different numbers of visits and different grooming behaviors and quantified pollen donor diversity on the pollinator’s body. We also evaluated the effect of grooming intensity and pollen removal or mixing on both pollen delivery and on the diversity of pollen donors received by stigmas.

## MODEL AND SIMULATIONS

### Model

We build agent-based models to simulate the deposition of individual pollen grains on the pollen vector’s body after sequential visits, the grooming behavior and effects of grooming on reproductive success of plants. Agent-based models are computational models that explore how individuals interact with each other and with their environment (Railsback and Grimm 2019). In this model, the agents are individual pollen grains, and the space corresponds to the vector’s body where the pollen grains are placed, mixed, removed or delivered to stigmas. In this study, we assume that pollen vector is a bee. The model components and process are described below.

The ABM has two components *i*.*e*. the flowers and the pollen vector. In this model, each plant bears a single bisexual flower with a fixed position of anther and stigma relatively to the vectors body. The population has a total of *f* flowers each one with different anther and stigma positions. Further floral traits are the following: mean number *r* of pollen grains released by anthers per visit, size α of the pollen patch, which affects the distribution of released pollen grains on bee’s body, and maximum number σ of pollen grains supported by stigma (Fig. S1). The vector’s traits used in our model describe a 3-dimensional space where pollen grains can be placed. Therefore, such traits are width (w) and height (h) and depth (d), an in our case w=h (Fig. S2). In our study, each individual bee can touch specific regions of its body, performing grooming behavior between sequences of visits. During grooming behavior, the bee either mixes the pollen grains from their original placement site to new sites on its body (mixing) or removes the pollen grains from their body (removal). We created two possible grooming behaviors: random (gr), in which the bee mixes and removes pollen randomly on its entire body; and ventral (gv), in which the bee mixes or removes pollen grains on the ventral thorax and abdomen. According to the arrangement of floral parts, the bee interacts with the flower following sequential processes during its visit: (I) pollen delivery to stigmas, (II) pollen placement by anthers and (III) grooming (Figure 1). To parameterize the model, such as the size of the bee’s body, amount of pollen released and amount of pollen received by the stigma, we used *Solanum lycocarpum* A. St. Hil (Solanaceae) and *Epicharis* Klug, 1807 (Apidae, Centridini). In this system, the anther places pollen in the center of the bee’s body and the stigma removes pollen from the same bee body’s region. The values used in the parameterization are based on previous investigation of this plant species floral traits and its pollinators (Marcelo et al. 2021) (Table 1).

**Table 1.**
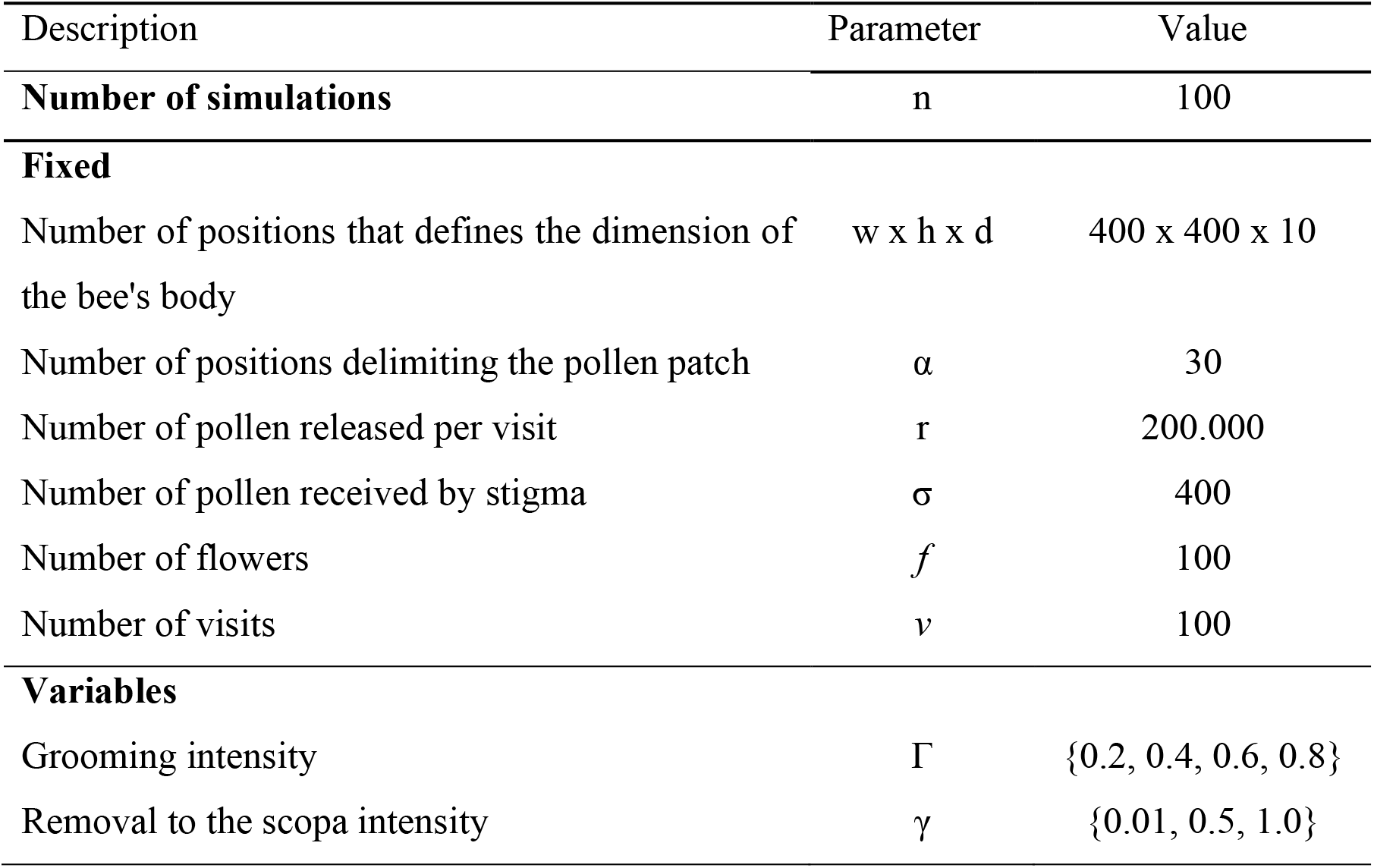
Description of parameter standard values used in the model parameterization, based on data from *Solanum lycocarpum* A. St. Hil (Solanaceae) and *Epicharis* sp. Klug, 1807 (Apidae, Centridini) (Marcelo et al. 2021). Some parameters are fixed: dimension of the bee’s body (which is height x width x depth, totaling 1,600,000 available sites for the pollen grains to be placed); pollen patch average position where pollen can be placed on the bee’s body); mean number of pollen grains released by anthers per visit; mean number of pollen grains delivered to stigmas; number of flowers created for floral visits and number of visits performed by bees. Other parameters were changed during the simulations: grooming intensity (proportion of sites on the body where the bee touches) and removal to the scopa intensity (proportion of sites the bee touches that will be removed to the scopa).

**Figure 1.**
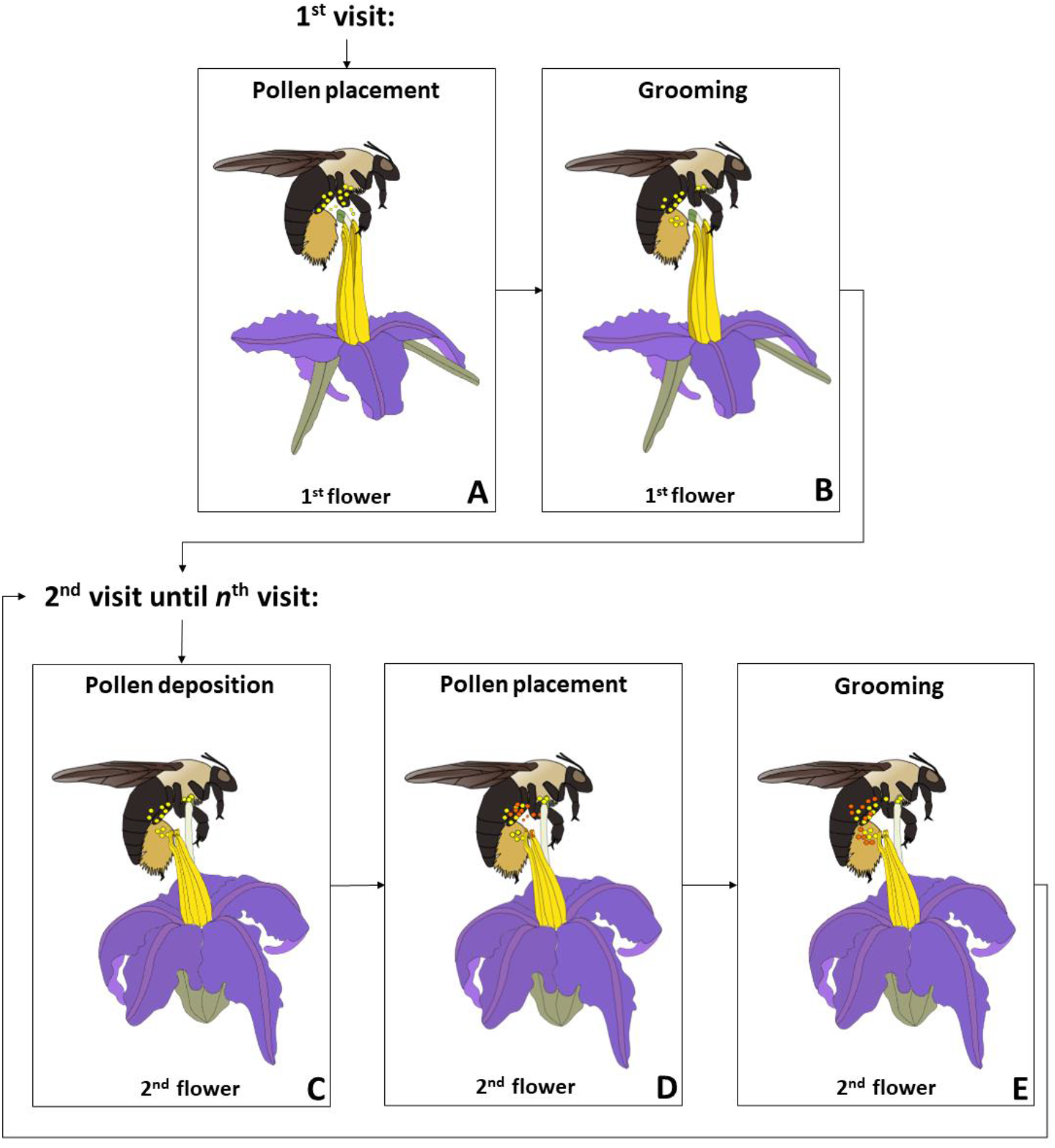
The rationale of the spatially explicit agent-based model used in this study to understand how pollen layering and grooming behavior in sequential visits may alter pollen landscape as well as pollen delivery. In the first flower visit, the bee’s body is clean since there was no previous pollen placement. Therefore, pollen deposition does not occur on the stigma of the first flower. Thus, in this first visit, only pollen placement by the anthers on the bee’s body can occur (A). Subsequently, the bee does the grooming behavior (B). In the second visited flower of the sequence, the bee already has pollen grains from the first flower visited. Therefore, the first action is pollen deposition, that is, the stigma of the second flower receives pollen from the bee’s body (C). Now, pollen grains from the anther of the second visited flower are placed on the bee (D), and, afterwards, the bee removes and mixes pollen in the grooming behavior (E). Such actions are repeated in the third visit and so on until the total number of visits – *v*, is achieved.

To understand the effect of grooming behavior on pollen landscape and plant reproduction, we performed 100 simulations combining different values of grooming and removal to the scopa intensities in bees with different grooming behaviors. Grooming intensity (Γ) represents the fraction of sites on the body that the bee touches, and we analyzed Γ= {0.2, 0.4, 0.6 or 0.8}. Removal to the scopa intensity is the fraction of the touched sites from which pollen will be displaced to the scopa. The fraction that is not displaced to the scopa will be relocated in another place of bee’s body γ = {0.01, 0.5, or 1}. Therefore, an extremely low value of removal to the scopa intensity, such as γ = 0.01, means that the bee is virtually mixing the pollen of all sites it touches. On the other hand, a high value of removal to the scopa intensity, as such the maximum γ = 1, means that all touched pollen grains will be removed from bee’s body and relocated in their scopa, and no mixing occurs.

### Simulations

The simulation starts with the creation of a sequence of different flowers (Fig. S3). The order of floral visit is randomly defined (without replacement). In our simulations each flower receives a single visit. The bee performs a total of *v* visits according to this sequence. Each flower has a respective stigma and anther positions (width and height) that correspond to the place where they touch the bee body. These positions are defined from a normal distribution with average on two hundred and variance of fifteen.

Then, the following processes happen (see Figure 1):

I. – Pollen delivery: previous pollen placed on bee’s body is delivered to the stigma of the current visited flower. The number and the identity of delivered pollen grains depends on both the region where the stigma touches and the availability of pollen grains in such region.
II. – Pollen placement: pollen grains from the anthers are placed on the bee’s body according to the position of the anther, the pollen patch (Fig. S4) and the amount of pollen released by anthers (Fig. S5). Each pollen placed has a position and layer on the bee’s body, which are labeled by their identity. If a specific position of the bee’s body is already fully occupied by pollen grains, pollen is lost during pollen placement.
III. – Grooming: bee grooming behavior is defined by the intensity Γ, which gives the proportion of sites that the bee can touch and the removal to the scopa value γ which gives the proportion of touched of sites from where pollen will be removed to the scopa. Pollen grains removed from the bee’s body to scopa will be unable to reach the stigma. The remaining pollen grains in touched sites are replaced in new sites of bee’s body due to mixing. The positions where the bee touches to remove and mix pollen grains are determined according to the grooming behavior (random grooming or thorax-ventral grooming).

### Analyses

We evaluated the spatial distribution of the total number of pollen grains and the number of pollen donors on the bee’s body simulating 1, 10, 50 and 100 visits by bees without grooming, and with random and ventral grooming. The diversity of pollen donors (Shannon-index) in each layer of the bee’s body was also quantified after 100 visits, according to different intensities of grooming and scopa values considering bees with random or ventral grooming. Finally, we evaluated pollen delivered by the first visited flower on the stigmas of the next 99 flowers as well the diversity (Shannon index) of pollen donors received by their stigmas after 100 visits considering the same intensities of grooming and removal to the scopa values, as well as each type of grooming behavior (random and ventral grooming). We performed 100 runs for each simulated bee.

All simulations were done in R version 4.0.2 (R Development Core Team 2020). The code is available at: (Code). We used the following packages: vegan (Oksanen et al. 2020); Rcpp (Eddelbuettel et al. 2022); dplyr (Wickham et al. 2019); plyr (Wickham 2020); purr (Henry and Wickham 2020); reshape2 (Wickham 2007); gsubfn (Grothendieck 2018); ggpubr (Kassambara 2020); plotrix (Lemon 2006); plot.matrix (Klinke and Chevalier 2022); ggplot2 (Wickham 2016); viridis (Garnier et al. 2021) and wesanderson (Karthik et al. 2018).

## RESULTS

### Distribution of pollen on the bee’s body

The sequential number of visits and the grooming behavior affected the spatial distribution of pollen grains on the bee’s body (Fig. 2). The total number of pollen grains increased with the number of visits in bees with or without grooming behavior. Specifically, pollen grains accumulated on the ventral regions of the thorax and abdomen in bees without grooming behavior, quickly filling the available sites for pollen placing. On the other hand, pollen grains were redistributed on the bee’s body never saturating the available sites when bees performed random or ventral grooming behavior. However, the distribution of the total number of pollen grains was different among bees with different grooming behaviors. Pollen grains accumulated on the lateral sides and of the grooming ventral region of the abdomen in the bee body performing ventral grooming behavior.

**Figure 2.**
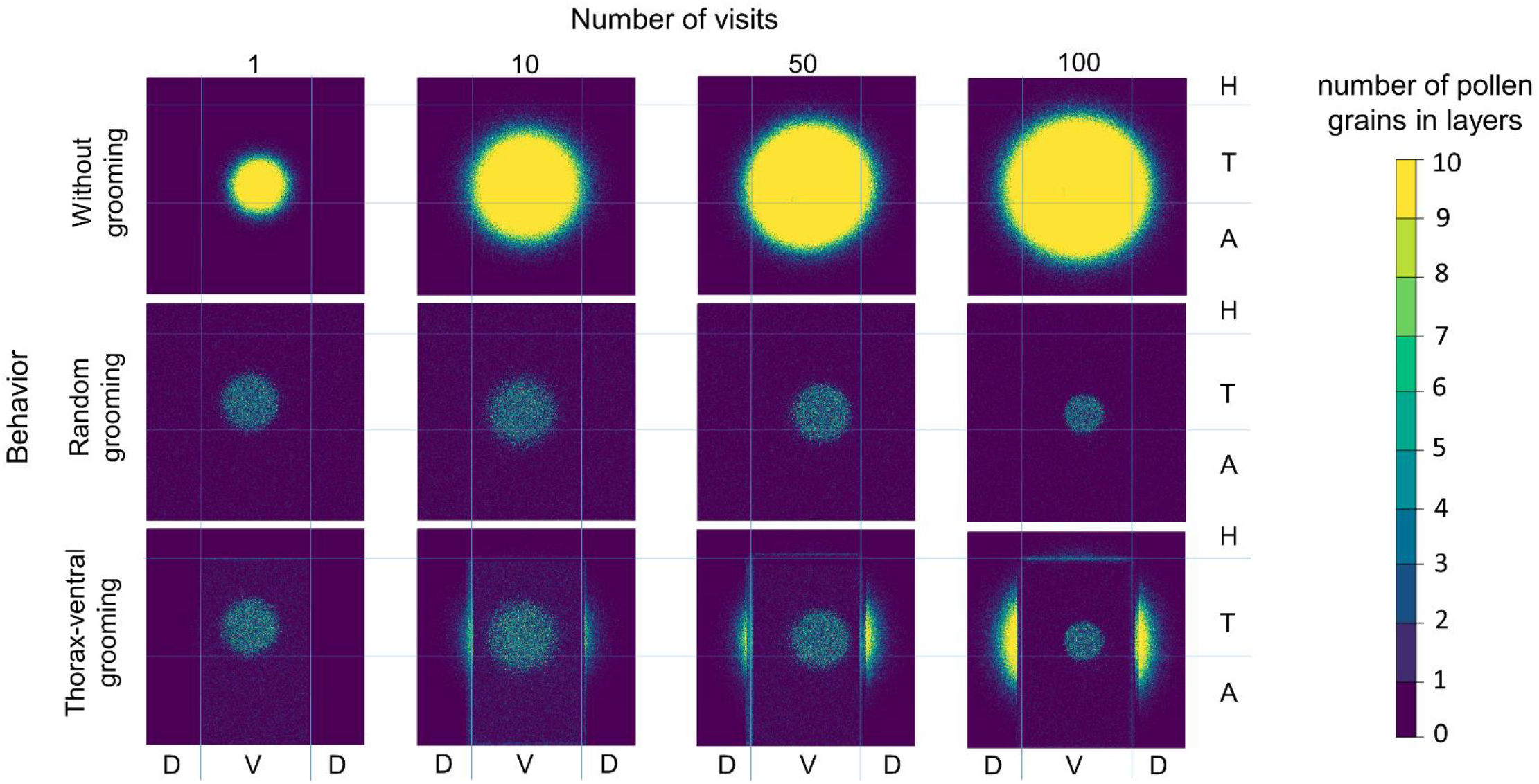
The total number of pollen grains accumulated on the bee’s body after different numbers of visits (1, 10, 50, and 100) and according to different grooming behaviors (without grooming, random grooming, and ventral grooming) in a single iteration. Blue lines indicate the boundaries of the bee’s body regions: horizontal lines delimit the head (H), thorax (T), and abdomen (A) regions while vertical lines delimit the ventral (V) and dorsal (D) regions. The yellow color indicates the highest number of pollen grains.

Sequential visits and grooming behavior also affected the spatial distribution of pollen grains from different donors on the bee’s body (Fig. 3). The central regions of the pollen patch deposited on the bee without grooming behavior were less rich in pollen donors than its periphery. Differently from the bee with random grooming behavior, the lateral sides of the bee performing ventral grooming behavior presented the highest number of different pollen donors.

**Figure 3.**
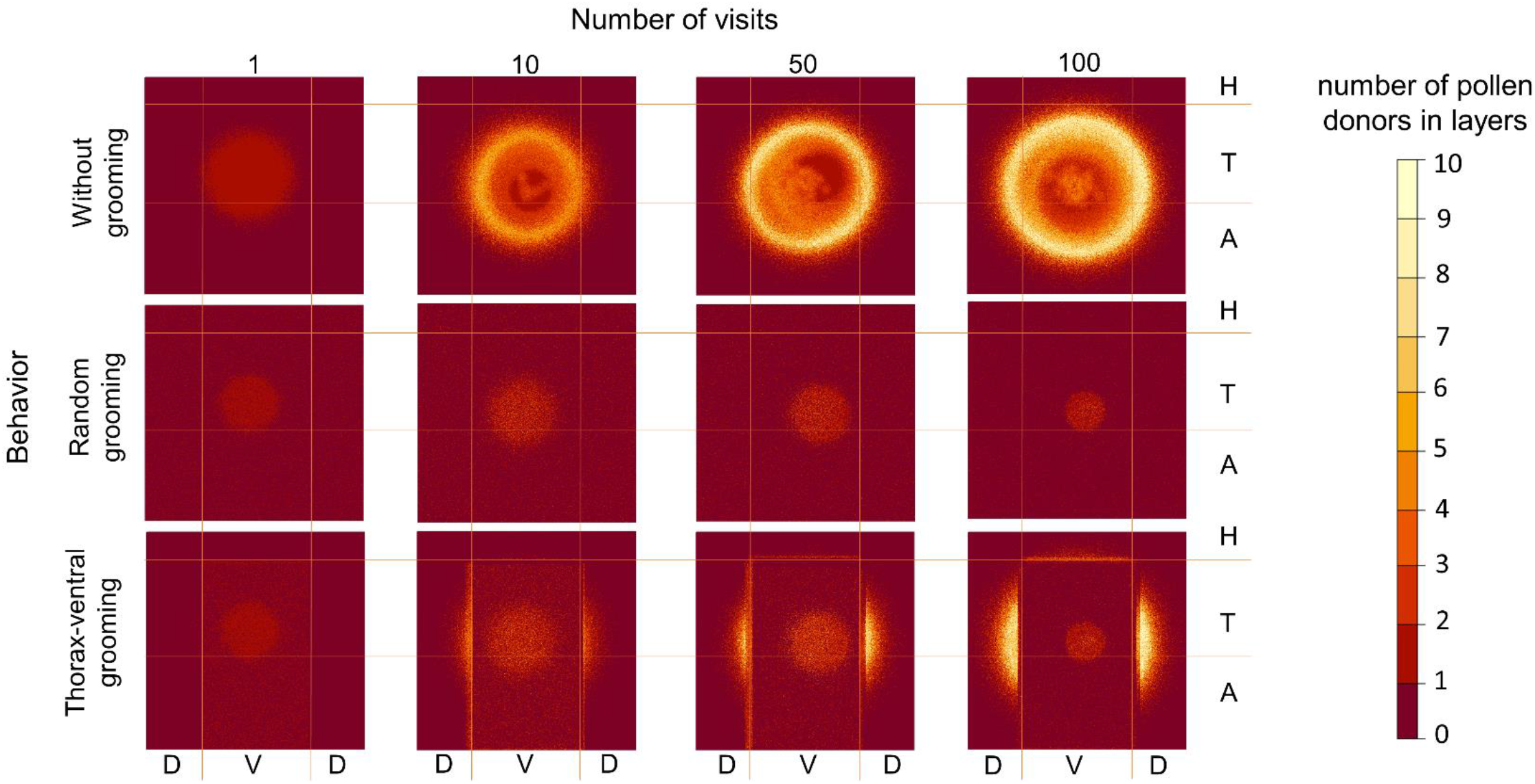
Number of different pollen grain donors on the bee’s body after different numbers of visits (1, 10, 50, and 100) and according to different grooming behaviors (without grooming, random grooming, and ventral grooming) in a single iteration. Yellow lines indicate the boundaries of the bee’s body regions: horizontal lines delimit the head (H), thorax (T), and abdomen (A) regions while vertical lines delimit the ventral (V) and dorsal (D) regions. The light ivory color indicates the highest number of different pollen donors.

### Pollen layers

The diversity of pollen donors on the bees’ body layers is affected by both the intensity of grooming behavior and the proportion of pollen grains moved to their scopa. In general, bees that mix almost all or half of the touched pollen grains in their venter (i.e., bees with low and intermediate values of scopa, respectively) bear higher pollen diversity than bees that move all touched pollen grains to scopa. However, the diversity of pollen grains is not evenly distributed across pollen layers. After 100 visits, the highest values of pollen diversity are found in intermediate layers, especially in bees with intermediate scopa values (Fig. 4A – C). In the high grooming intensity scenario, the inner layers present the greater pollen diversity when the bee removes all or intermediate amounts of pollen grains to scopa (Fig. 4D). The diversity of pollen grains across layers is similar for bees with random grooming behavior (Fig. S6).

**Figure 4.**
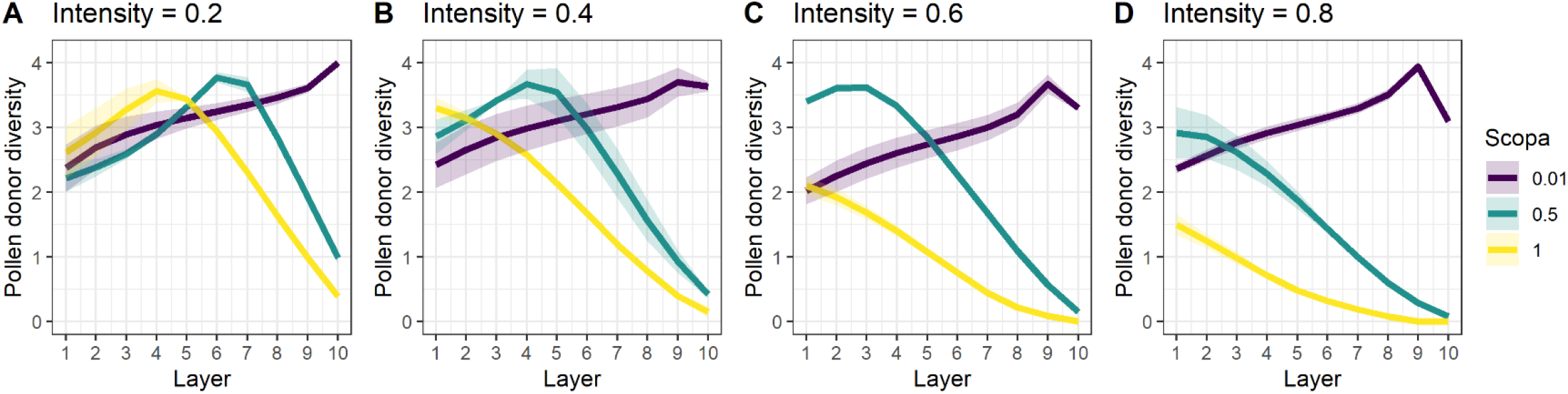
Diversity (Shannon index values) of pollen grains on the layers of the bee’s body according to different intensities of grooming and scopa values in ventral grooming bees after 100 floral visits. On the x-axis, the higher the layer number, the outermost the layer. Line colors indicate scope intensities, with purple 0.01, navy-blue 0.5 and yellow 1.0. Number of iterations = 100.

### Grooming effect on plant reproduction

Although the cumulative curve of pollen delivered by the first visited flower increased in sequential visits, total pollen delivered decreases with grooming intensity in bees with different behavior (Fig. 5A - D). Furthermore, the cumulative curve of pollen delivered behaves differently in bees with different grooming behaviors. The cumulative curve of pollen delivery saturates faster in bees that move all pollen grains to their scopa as compared with bees that mix pollen grains on their bodies, especially in higher values of grooming intensity. Higher values of pollen delivery were achieved by bees with low scopa values, indicating that pollen mixing increases the total number of pollen grains delivered as well as the number of pollinated flowers.

**Figure 5.**
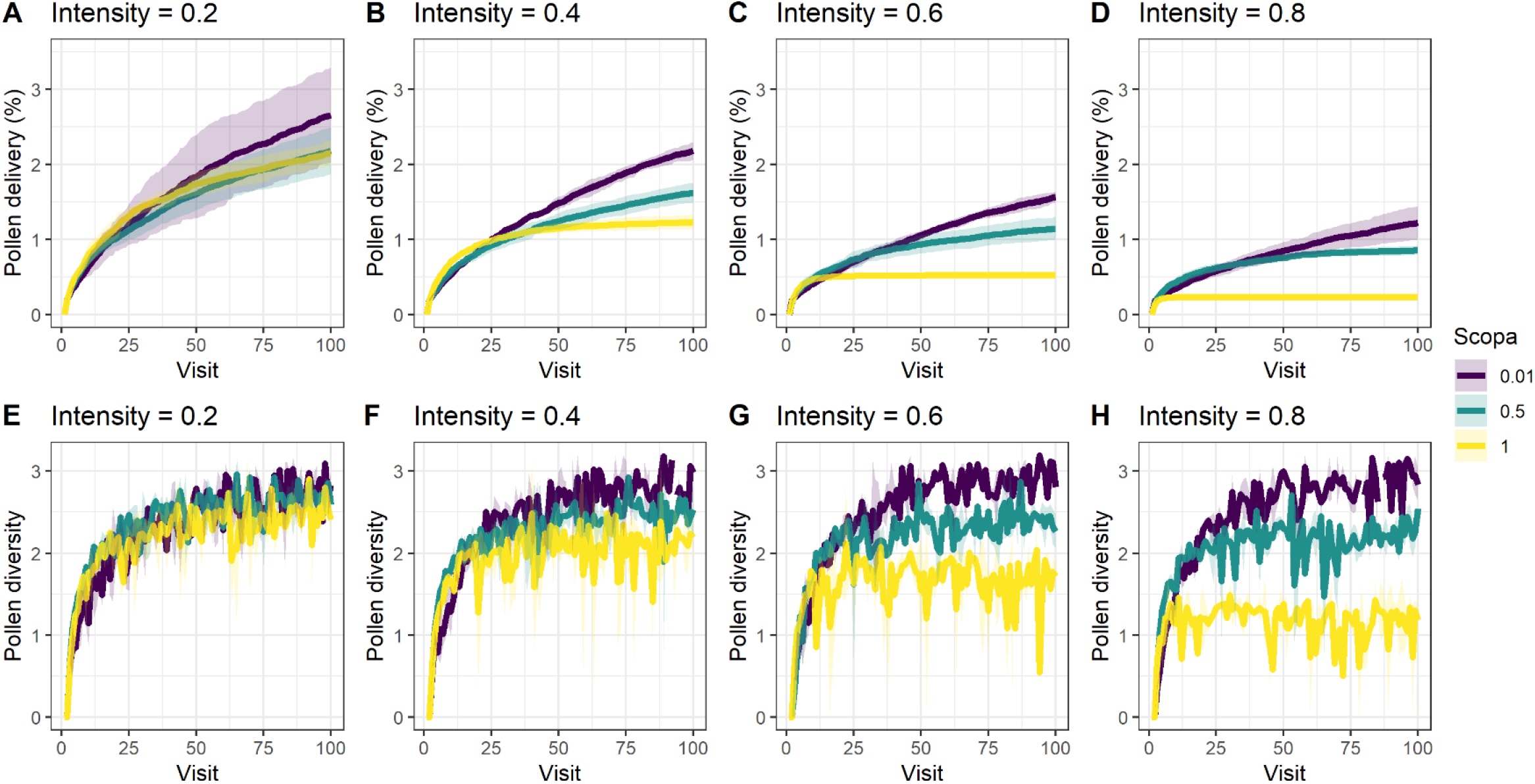
Effect of grooming intensity and removal to the scopa on fitness of the first visited flower, measured by the proportion of released pollen grains from the first flower that is delivered to stigmas of conspecific flowers (A-D). And effect of grooming intensity and removal to the scopa on the diversity (Shannon index) of pollen grains received by stigmas in sequential visits (E-H). Line colors indicate scope intensities, with purple 0.01, navy-blue 0.5 and yellow 1.0. Number of iterations = 100.

The diversity of pollen deposited on stigmas increased throughout sequential floral visits in all intensity scenarios and all simulated bees (Fig. 5E – H). However, in high values of grooming intensity, stigmas receive higher diversity of pollen grains from bees with low to intermediate values of scopa as compared with bees that move all pollen grains to their scopa in sequential visits. Similar results for pollen delivery and diversity of pollen deposited in stigmas in sequential visits were found in simulated bees with random grooming (Fig. S7). In general, the results are not drastically affected by the setting of the parameter values (see the parameter space analysis in Fig. S8 – S11).

## DISCUSSION

Our simulations illustrate that grooming behavior has the potential to significantly restructure the spatial distribution of the pollen grains on the bee’s body, altering both the horizontal and vertical pollen landscape and, consequently, affecting flower reproductive success. Specifically, pollen accumulated on the unreachable lateral regions of the grooming bee’s body. In addition, pollen mixing due to bee’s grooming behavior results in a peak of maximum pollen diversity at intermediate layers. Thus, by altering the pollen landscape, grooming behavior has the potential to influence the number of pollen grains delivered to other flowers as well as the diversity of pollen grains deposited on stigmas.

### The effect of grooming on pollen landscape

During their foraging bout, bees sequentially visit flowers in search of pollen and/or nectar (Westerkamp 1996). The sequential pollen placement from different flower donors may result in a pollen landscape where pollen grains from different donors and in different amounts are placed on the bee’s body (Minnaar et al. 2019). Our study shows that, by removing or mixing pollen grains, heterogeneous grooming behavior has the potential to restructure this landscape. With the constant removal of pollen from the central regions of their body, the available sites in this grooming region never saturate, allowing the reception of pollen grains from new donors in each floral visit. Consequently, the central ventral region, where anthers and stigmas interact with pollen grains in our model, is less rich in pollen donors while there is an accumulation of pollen grains from different donors in the lateral regions of the bee’s body. Sites where bees do not touch pollen grains are called safe sites. Such sites have been reported for natural bees and may create hot and cold spots of outcrossed pollen (Koch et al. 2017; Tong and Huang 2018; Minnaar and Anderson 2021). Although we did not directly address the impact of hot and cold spots on the evolution of anthers and stigmas position on the bee’s body in our study, it is expected that both would evolve to positions that favor the contact with such safe sites, especially in flowers which pollen grains are a feeding resource for bees and a reproductive resource for plants. In fact, during the evolutionary history of pollen flowers, the position of anthers and stigmas has recurrently evolved to such positions, in some cases resulting in enantiostyly (Vallejo-Marín et al. 2010; Barrett 2021; Melo et al. 2021).

The sequential placement of pollen grains in similar regions of the bee’s body potentially creates pollen layers (Lertzman and Gass 1983). However, the diversity distribution along the vertical dimension of the pollen landscape can be drastically altered by bee grooming behavior. In bees with intermediate values of removal to the scopa, the constant removal and mixing of pollen grains from superficial layers generate a pattern of high pollen diversity in intermediate layers. Higher diversity values in regions with neither too rare nor too frequent disturbance have been hypothesized and demonstrated in several ecological systems, especially at the community level (Moi et al. 2020). In such dynamic and non-equilibrium systems, diversity increases when a new patch of an unoccupied habitat is available until the access to resources becomes limited and lower competitors are displaced or even excluded. Some disturbance in this system will make new habitats available for colonization by reducing the abundance of higher competitors, consequently resetting the system to some earlier higher diverse stage (Osman 2015). Interestingly, our model predicts that the intermediate disturbance hypothesis (Connel 1978; Grime 1973) could be corroborated on a tiny scale (i.e., pollen grains on the bee’s body). The development of techniques to track the identity of individual pollen grains from several and different plants in different pollen layers might allow an empirical test of this prediction and a better comprehension of pollen landscapes on vectors bodies.

### The effect of grooming on the reproductive success of plants

Pollen grains produced in flowers pollinated by animal vectors often present pollenkitt, a sticky coat material that promotes pollen grains aggregation and helps in their transport (Pacini and Hesse 2005), potentially facilitating the formation of multiple pollen donor layers. In contrast to previous studies (e.g., Minnaar et al. 2019), our theoretical model assumes that the vertical dimension of the bee’s body is not infinite, and that pollen layering does not increase indefinitely with sequential visits. In this sense, first placed pollen grains may have a spatial competitive advantage as compared to pollen placed in further visits. However, this competitive advantage would decrease when vectors perform more intense pollen grooming with some level of mixing. In fact, grooming behavior generally mixes pollen on grooming vectors, like bees (Harder and Barrett 1996).

In natural systems (Harder & Thomson 1989, Gong & Huang 2014), as well as in our simulations, approximately 5% of the pollen grains placed on the vector’s body reach conspecifics stigmas. In simulations with some levels of mixing, both the total number of pollen grains and the number of different flowers pollinated (i.e., pollen carryover) were higher than in cases when the bees move all pollen grains to their scopa. Pollen carryover is an important component of male fitness and affects the extent of gene flow (Morris et al. 1994). If the success of crosses between near neighbors were lower than crosses between moderately spaced plants (Levin 1984), the increase in the pollen carryover would increase the reproductive success of pollen donors (Morris et al. 1995) and reduce the effects of geitonogamy in mass-flowering plants (Morris et al. 1994). Pollen carryover is reduced by the sequential pollen placement in non-grooming vectors (Price & Waser 1982). Previous computer simulations suggested that such a decrease may be explained by the effect of pollen dilution or layering (Lertzman 1981; Lertzman & Grass 1982). Our simulations suggest that, if there is some degree of pollen mixing during grooming behavior, pollen carryover may be increased by grooming vectors, possibly by restructuring the pollen layered landscape (but see Thomson 1986). As pollen grains remain viable for a long period (Pacini and Franchi 2020), the positive effect of grooming on pollen delivery may even be higher than what has been predicted in our theoretical model, since a single bee can perform more than 300 visits in a day (Raine and Chittka 2007).

Finally, pollen grains are not necessarily successful when they are deposited on stigmas. As the available space for pollen tube growth within styles and the total number of ovules are limited, post-pollination processes such as pollen tube competition and female choice should have direct reproductive implications (Moore and Pannel 2011). Our model predicted that the diversity of pollen deposited on stigmas by grooming vectors can reach twice as higher values as the diversity of pollen delivered by non-grooming vectors, especially in high intensities scenarios. High diverse pollen loads enhance the chances of receiving pollen from a high compatible donor, increasing the number of fertilized ovules, reducing seed and fruit abortion (Janzen 1977; Stephenson 1981; Sutherland 1987). The diversity of pollen donors on stigmas may also enhance the quantity and quality of the progeny through interactions among pollen donors because pollen tubes may grow faster in the presence of competitors (Lankinen and Skogsmyr 2002; Kron and Husband 2006). Therefore, if grooming behavior promotes some pollen mixing on the vector’s body, it may ultimately enhance reproductive performance via seed quantity and quality. Given the ubiquity of bee pollinators, we expect that grooming behavior could often directly affect the evolution of floral traits that affect pollen placement on the pollinator’s body.

## ACKNOWLEDGMENTS

This work was carried out with the support of the Coordenação de Aperfeiçoamento de Pessoal de Nível Superior (CAPES) - Finance Code 001 - and Conselho Nacional de Desenvolvimento Científico e Tecnológico (CNPq) - Process: 308107/2021-7. The authors thank the members of the Vallejo-Marín Lab for their suggestions in the text, especially Carlos E. P. Nunes. We thank Pamela Santana and Pedro Bergamo for our valuable discussions.

