## Supplementary material for "POLLINATOR GROOMING BEHAVIOR ALTERS POLLEN LANDSCAPES ON BEES’ BODIES AND INCREASES POLLEN CARRYOVER TO OTHER FLOWERS"

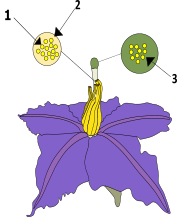

### **Figure S1.** Each flower created has a 1- mean number of pollen grains released by anthers per visit (r); 2- pollen patch (α) and 3-maximum number of pollen grains supported by stigma (σ).

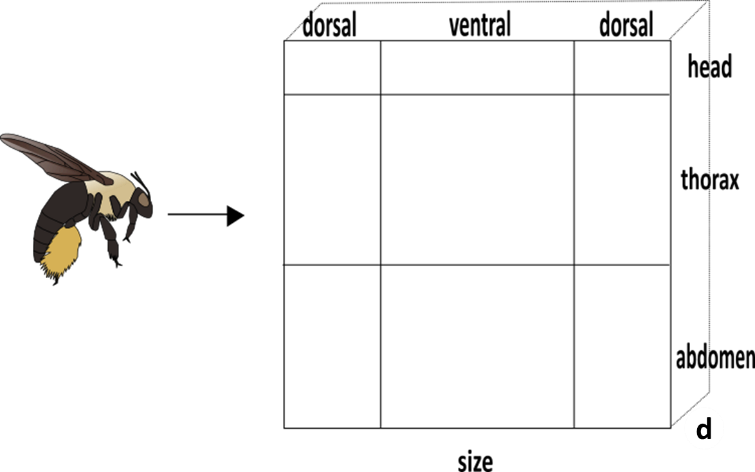

### **Figure S2.** The modeled bee has a size defined by its width (w), height (h) (in our case w=h) and depth (d). We also defined the bee´s body regions. Lines indicate the boundaries of the bee's body regions: the horizontal lines delimit the head, thorax and abdomen regions and the vertical lines delimit the ventral and dorsal regions.

flower 1

flower 2

flower 10

flower 9

flower 5

flower 3

flower 8

flower 6

flower 4

flower 7

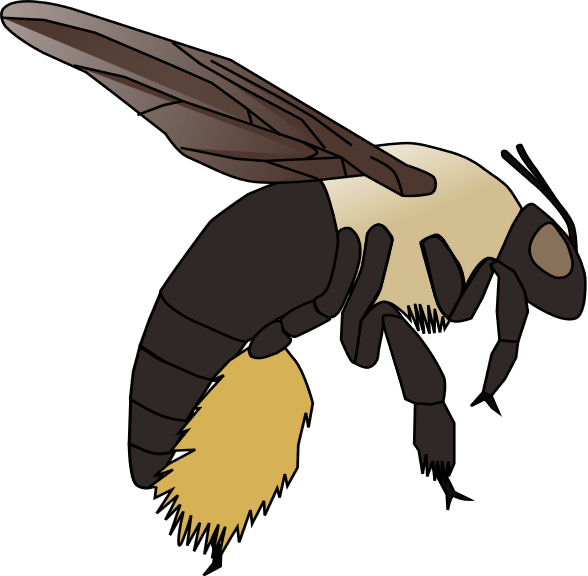

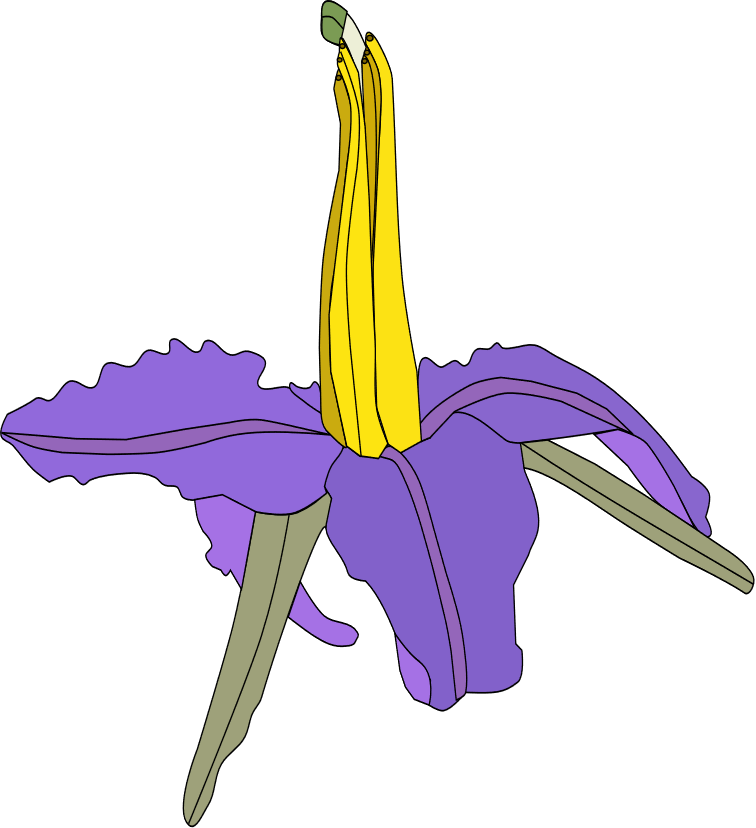

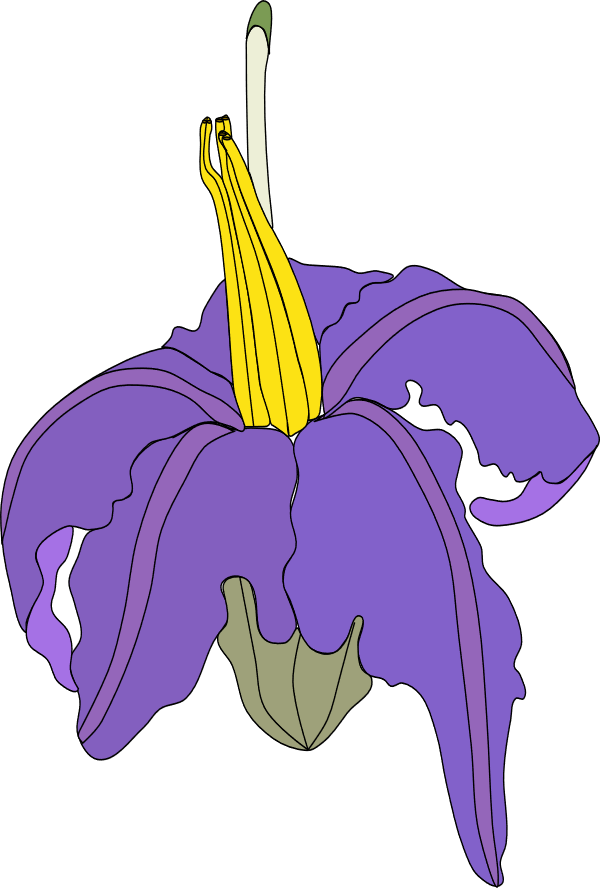

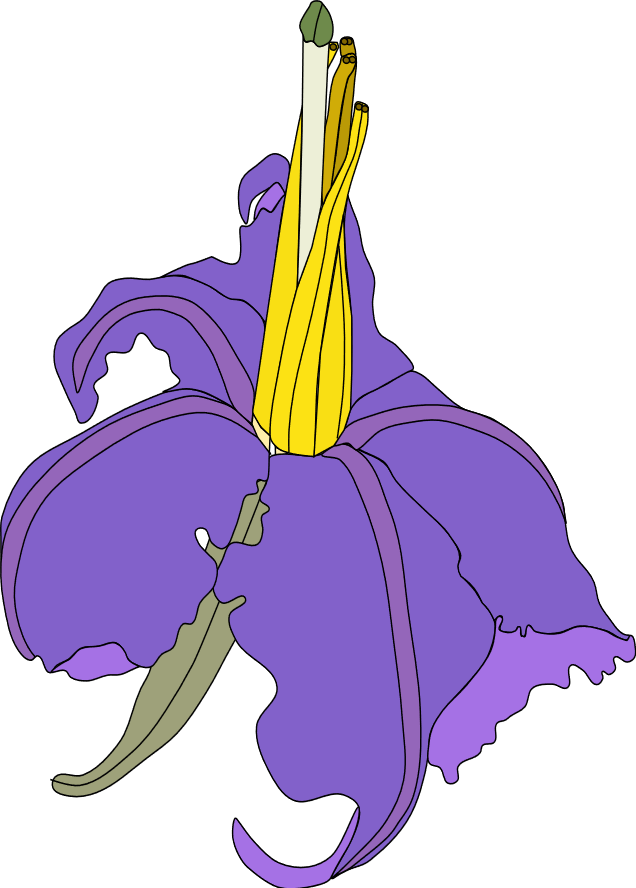

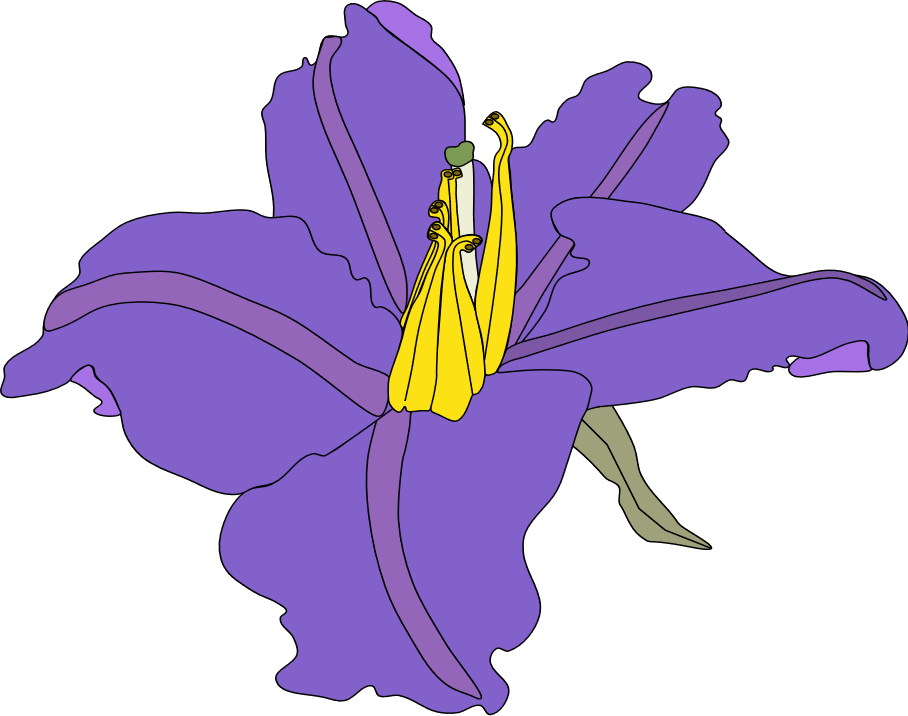

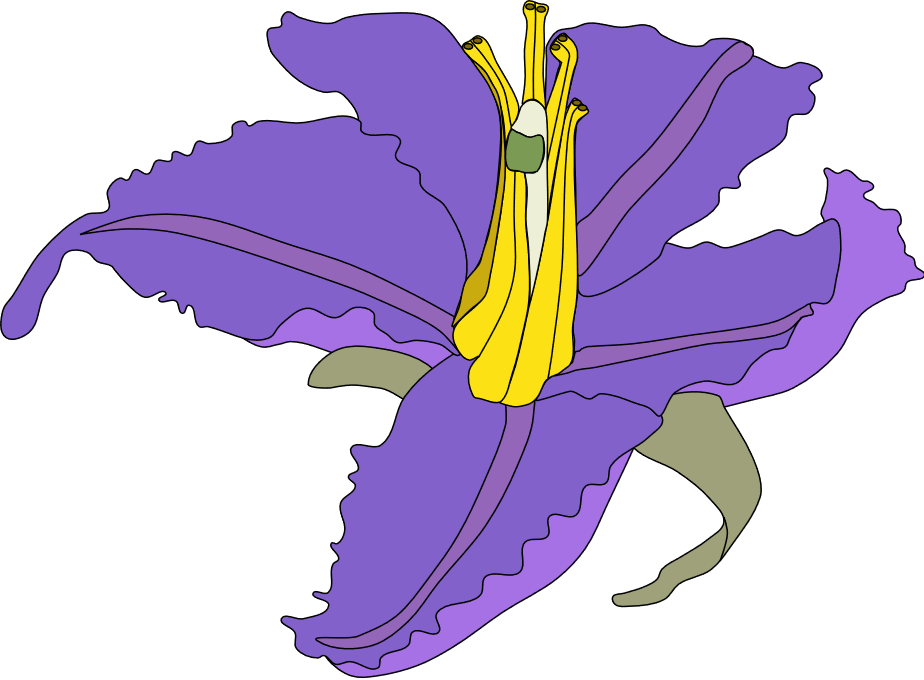

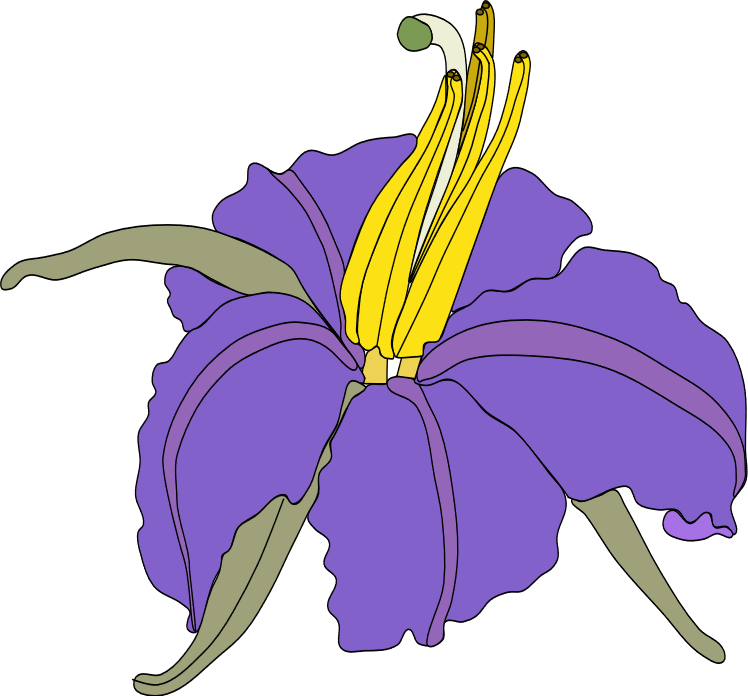

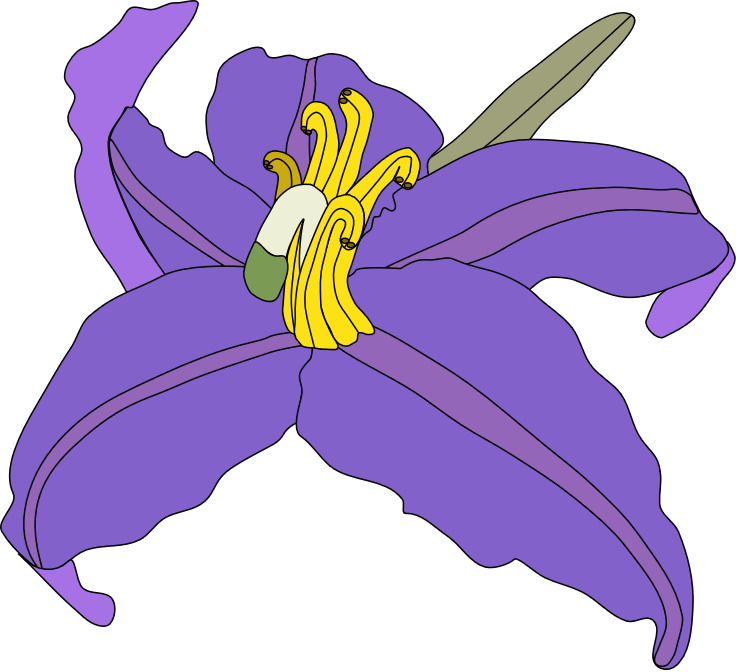

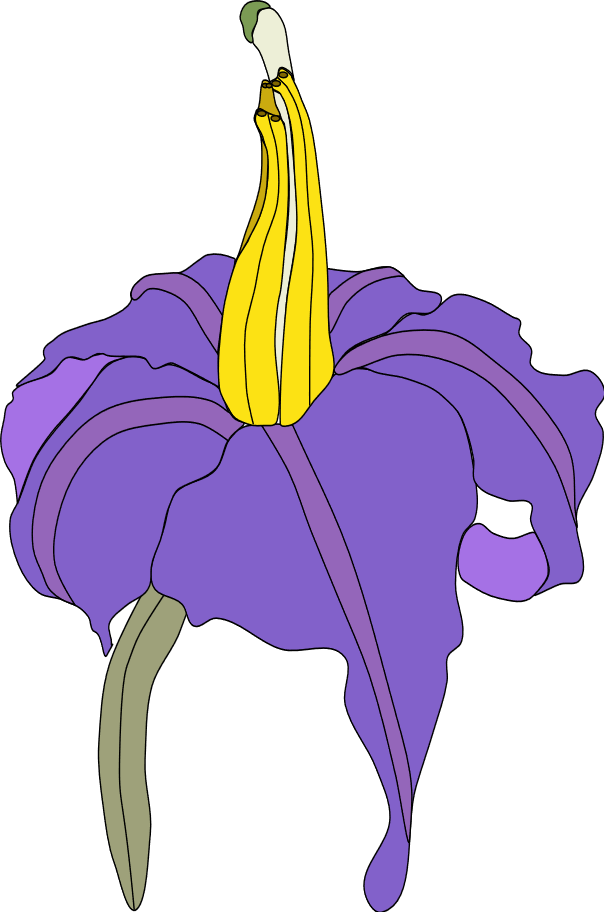

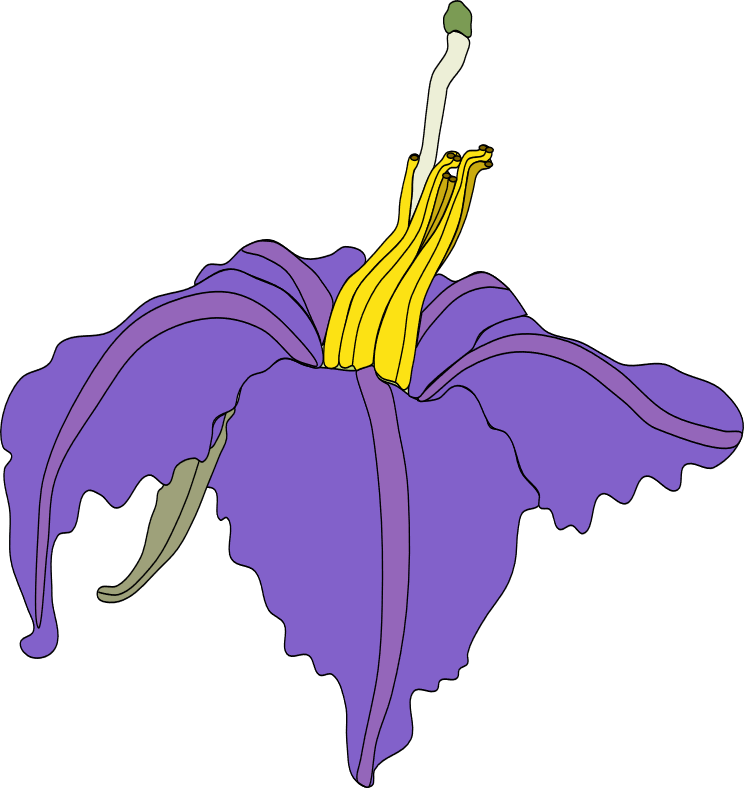

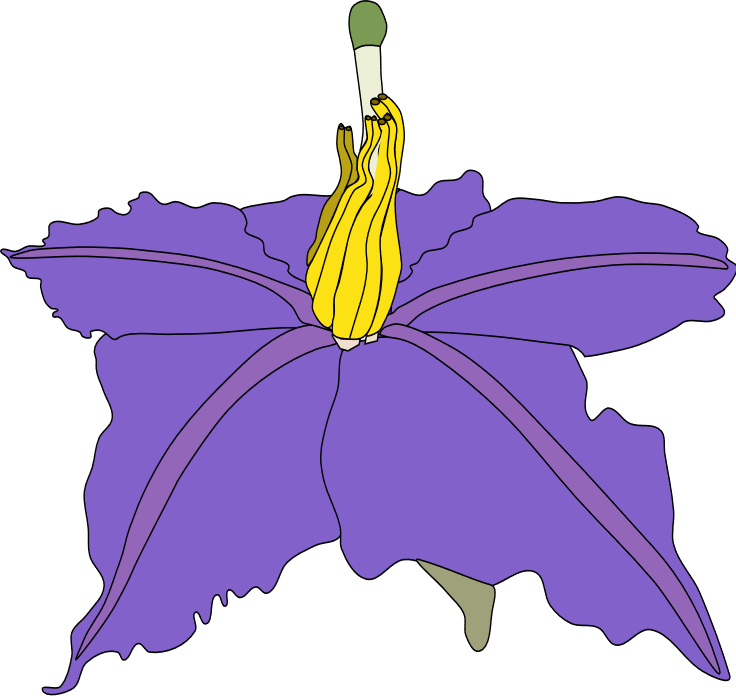

### **Figure S3.** Example of a random sequence of ten flowers created which the bee will delivery and collect pollen.

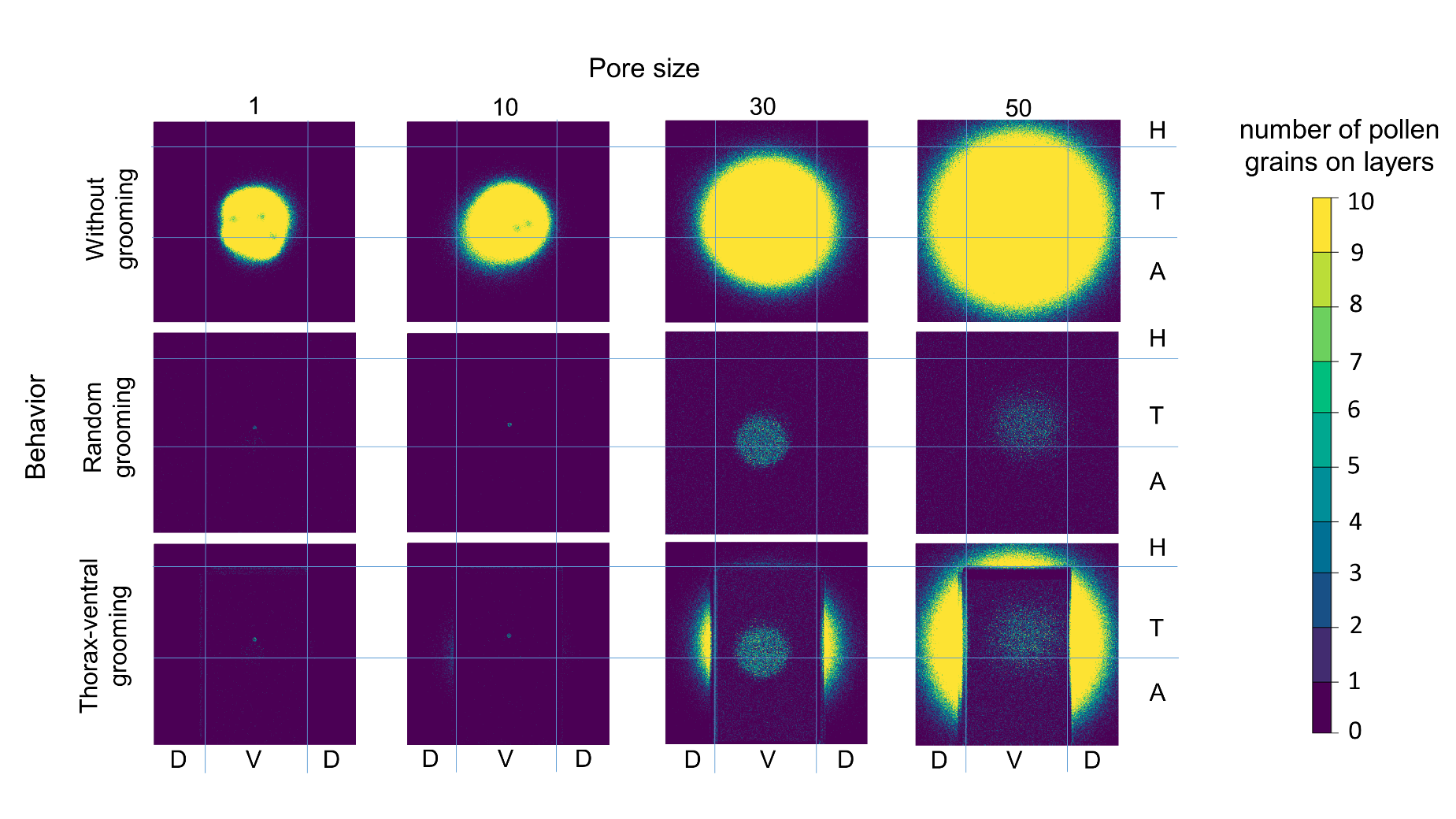

**Figure S4.** The total number of pollen grains accumulated on the bee´s body after one hundred visits in flowers with different pollen patch (1, 10, 30 and 50) and according to different grooming behaviors (without grooming, random grooming, and ventral grooming) in a single iteration. Blue lines indicate the boundaries of the bee's body regions: horizontal lines delimit the head (H), thorax (T), and abdomen (A) regions while vertical lines delimit the ventral (V) and dorsal (D) regions. The yellow color indicates the highest number of pollen grains.

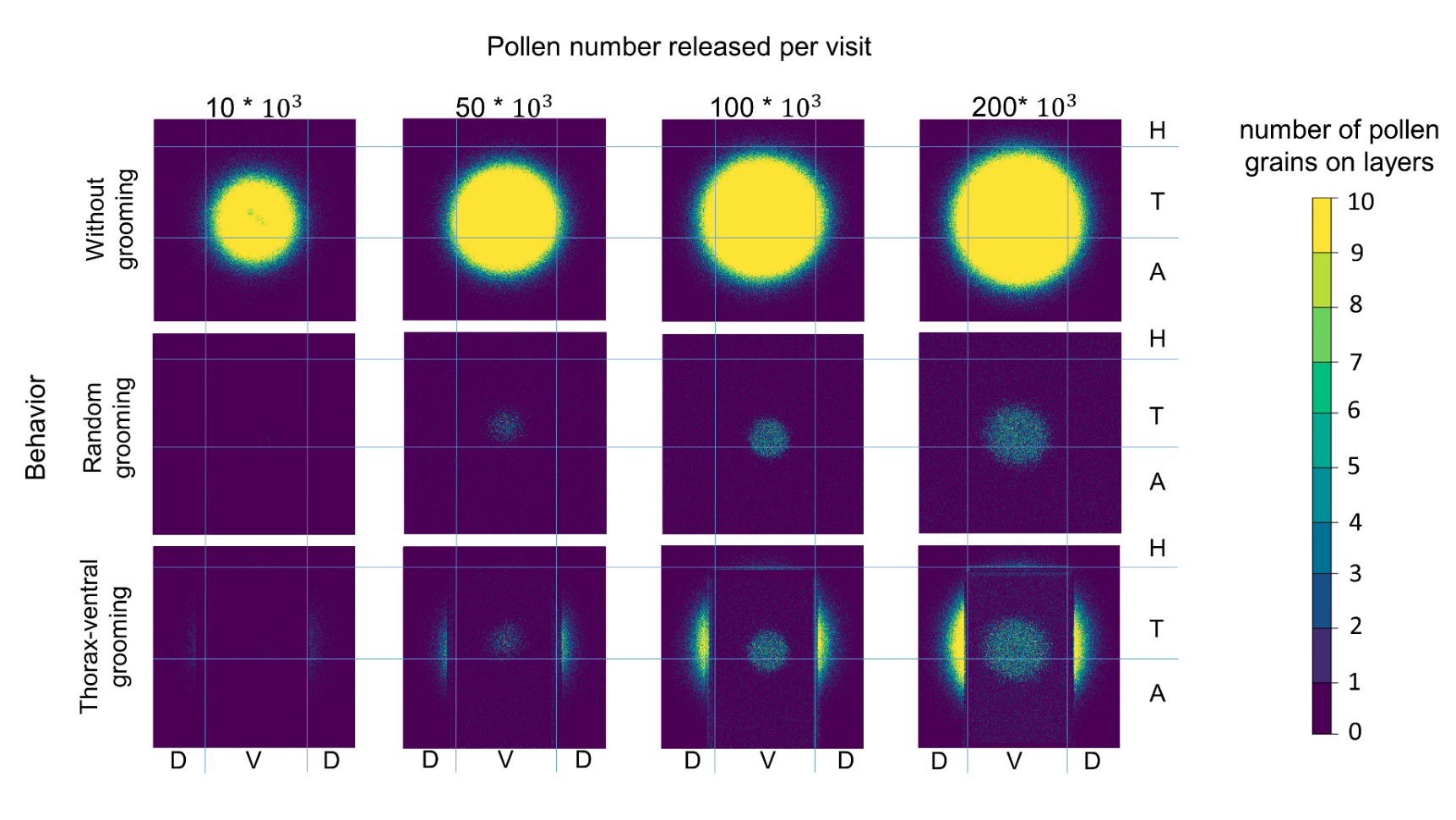

### **Figure S5.** The total number of pollen grains accumulated on the bee´s body after a hundred visits in flowers with different pollen numbers released per visit (10 x 10³, 50 x 10³, 100 x 10³ and 200 x 10³), and according to different grooming behaviors (without grooming, random grooming, and ventral grooming) in a single iteration. Blue lines indicate the boundaries of the bee's body regions: horizontal lines delimit the head (H), thorax (T), and abdomen (A) regions while vertical lines delimit the ventral (V) and dorsal (D) regions. The yellow color indicates the highest number of pollen grains.

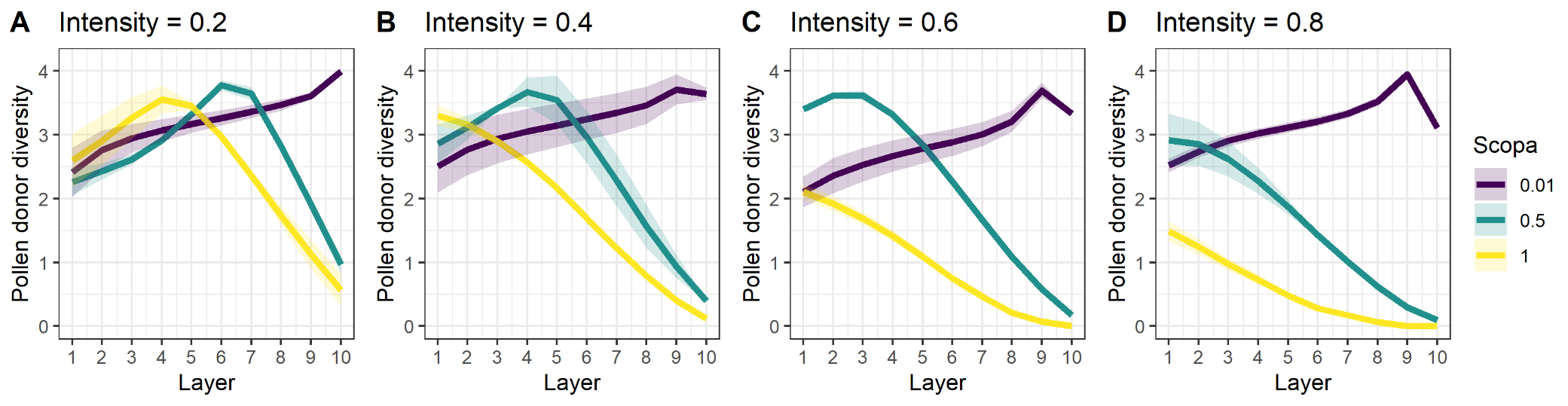

**Figure S6.** Diversity (Shannon index values) of pollen grains on the layers of the bee’s body according to different intensities of grooming and removal to the scopa values in random grooming bees after 100 floral visits. On the x-axis, the higher the layer number, the outermost the layer. Number of iterations = 100.

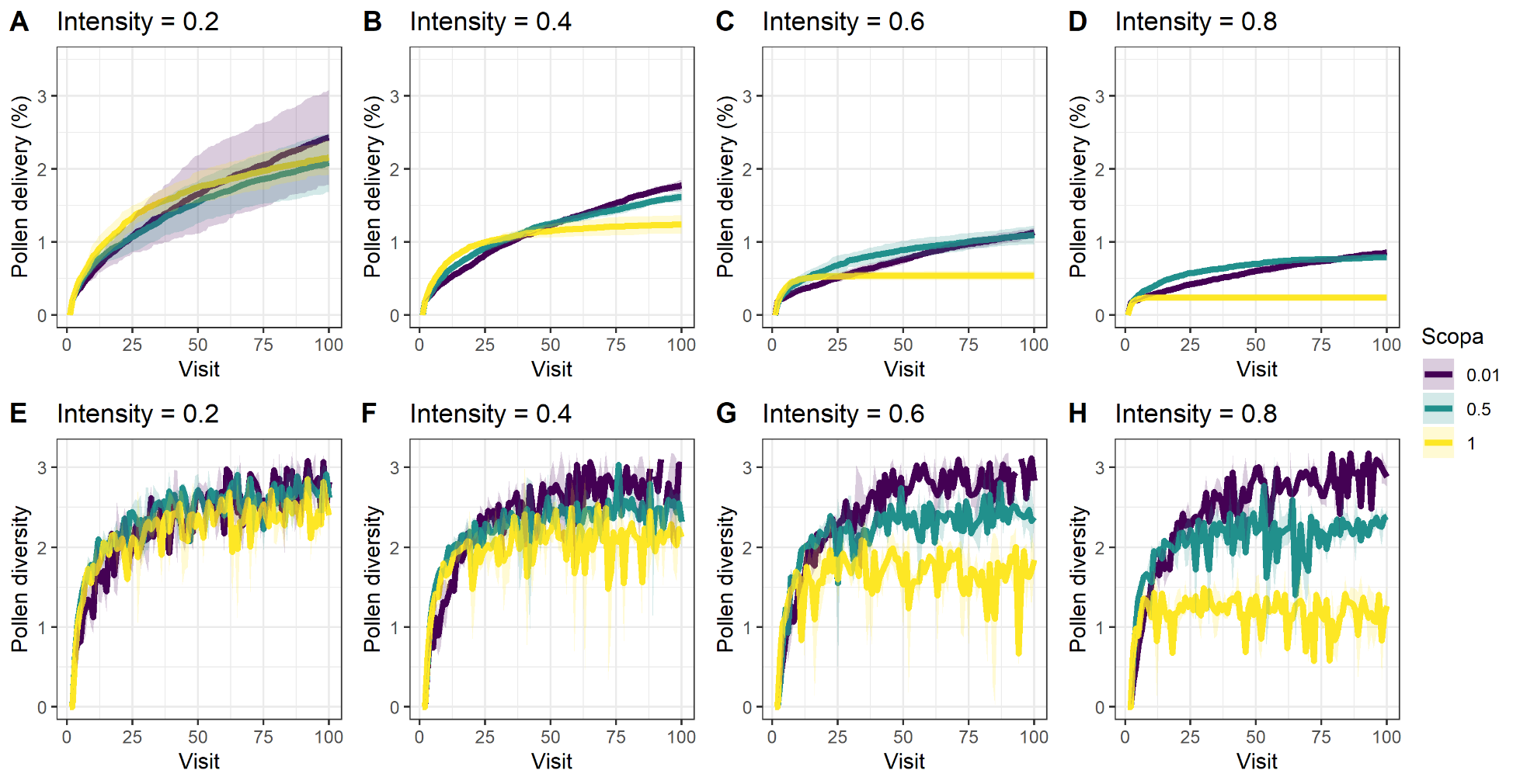

**Figure S7.** Effect of grooming intensity and removal to the scopa on the fitness of the first visited flower, measured by the proportion of released pollen grains from the first flower that is delivered to stigmas of conspecific flowers (A-D). And effect of grooming intensity and removal to the scopa on the diversity (Shannon index) of pollen grains received by stigmas in sequential visits (E-H). Number of iterations = 100.

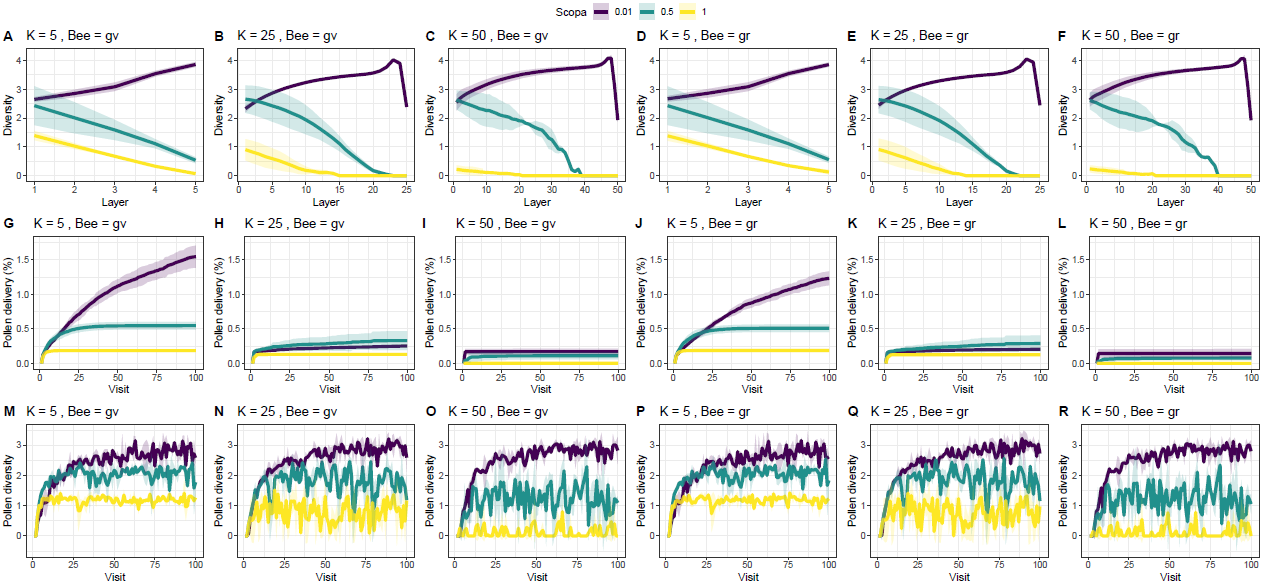

**Figure S8.** K parameter space analysis. k indicates the number of layers in the bee's body. Plots show (A-F) pollen diversity in bee body layers, (G-L) percentage of pollen from the first flower delivered to conspecific stigma, and (M-R) pollen diversity on stigmas in sequential visits, when k is 5 (A, G ,M, D, J ,P), 25 (B, H, N, E, K, Q) and 50 (C, I, O, F, L, R) and when behavior is thorax-ventral grooming (A, B, C, G, H, I, M, N, O) and random grooming (D, E, F, J, K, L, P, Q, R). Line colors indicate scope intensities, with purple 0.01, navy-blue 0.5 and yellow 1.0.

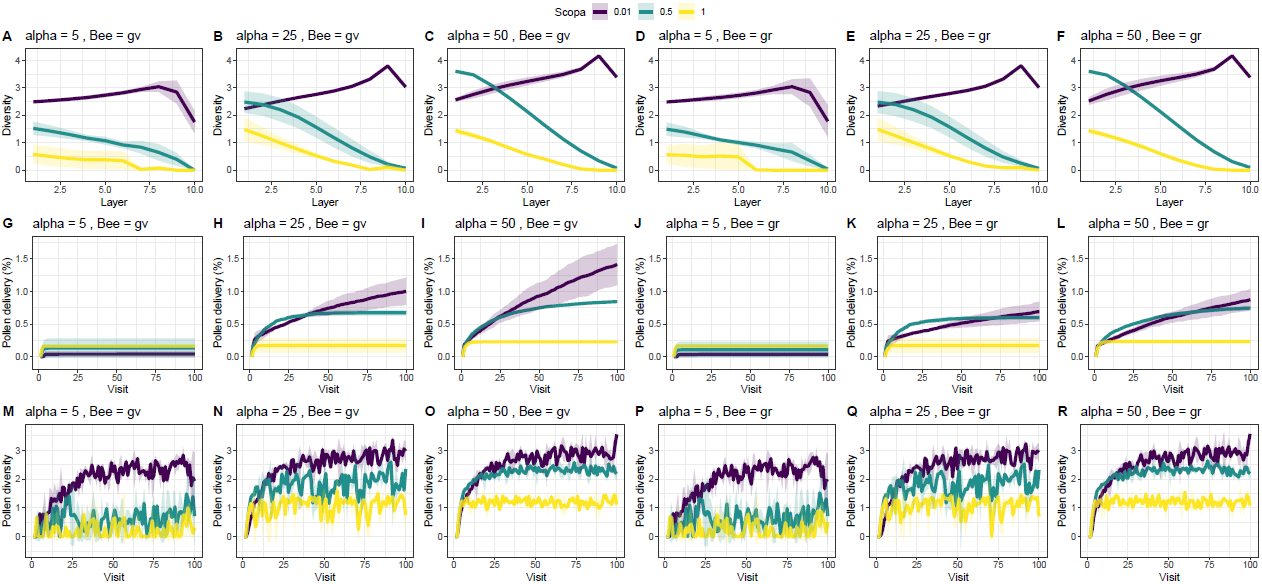

**Figure S9.** Alpha parameter space analysis (α). Alpha indicates the number of positions that delimit the pollen patch. Plots show (A-F) pollen diversity in bee body layers, (G-L) percentage of pollen from the first flower delivered to conspecific stigma, and (M-R) pollen diversity on stigmas in sequential visits, when alpha is 5 (A, G ,M, D, J ,P), 25 (B, H, N, E, K, Q) and 50 (C, I, O, F, L, R) and when behavior is thorax-ventral grooming (A, B, C, G, H, I, M, N, O) and random grooming (D, E, F, J, K, L, P, Q, R). Line colors indicate scope intensities, with purple 0.01, navy-blue 0.5 and yellow 1.0.

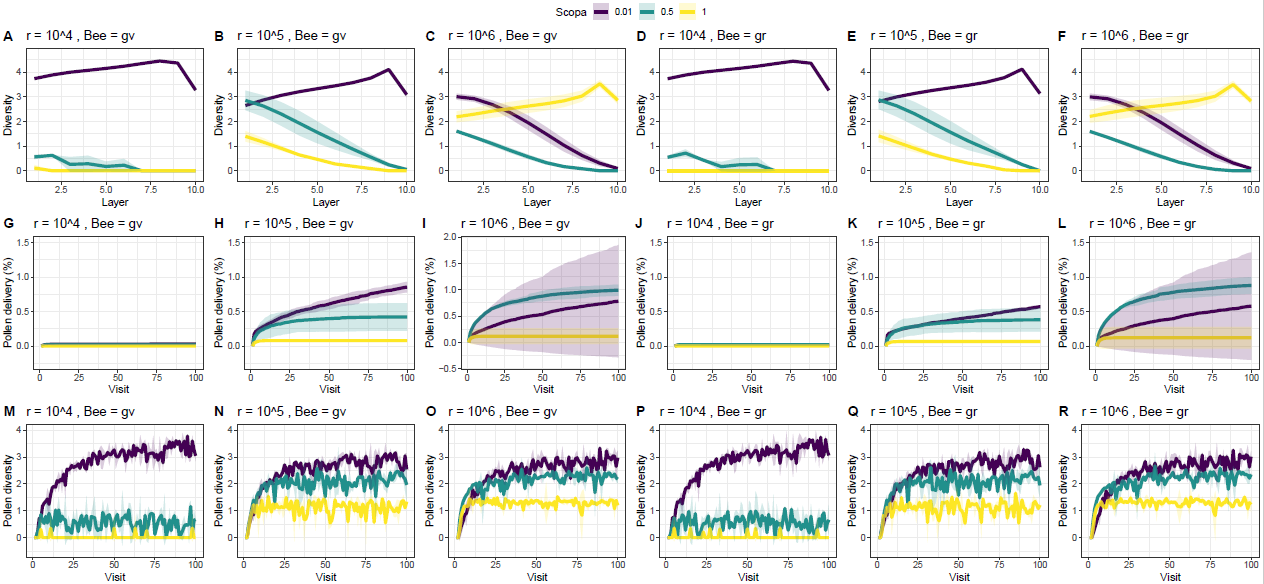

**Figure S10.** R parameter space analysis. r indicates the number of pollen released per visit. Plots show (A-F) pollen diversity in bee body layers, (G-L) percentage of pollen from the first flower delivered to conspecific stigma, and (M-R) pollen diversity on stigmas in sequential visits, when r is 10^4^ (A, G ,M, D, J ,P), 10^5^ (B, H, N, E, K, Q) and 10^6^ (C, I, O, F, L, R) and when behavior is thorax-ventral grooming (A, B, C, G, H, I, M, N, O) and random grooming (D, E, F, J, K, L, P, Q, R). Line colors indicate scope intensities, with purple 0.01, navy-blue 0.5 and yellow 1.0.

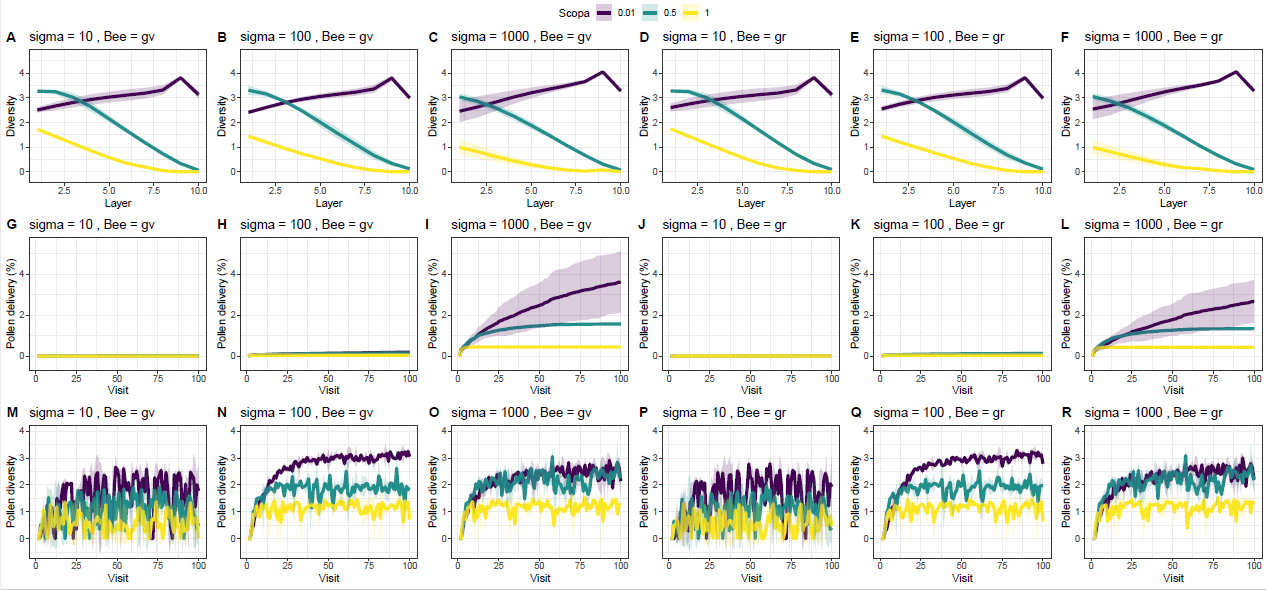

**Figure S11.** Sigma parameter space analysis (σ). Sigma indicates the number of pollen received by stigma. Plots show (A-F) pollen diversity in bee body layers, (G-L) percentage of pollen from the first flower delivered to conspecific stigma, and (M-R) pollen diversity on stigmas in sequential visits, when sigma is 10 (A, G ,M, D, J ,P), 100 (B, H, N, E, K, Q) and 1000 (C, I, O, F, L, R) and when behavior is thorax-ventral grooming (A, B, C, G, H, I, M, N, O) and random grooming (D, E, F, J, K, L, P, Q, R). Line colors indicate scope intensities, with purple 0.01, navy-blue 0.5 and yellow 1.0.
